## Supplementary figures and images for "Deep-learning and transfer learning identify new breast cancer survival subtypes from single-cell imaging data"

### Supplemental Figure 1

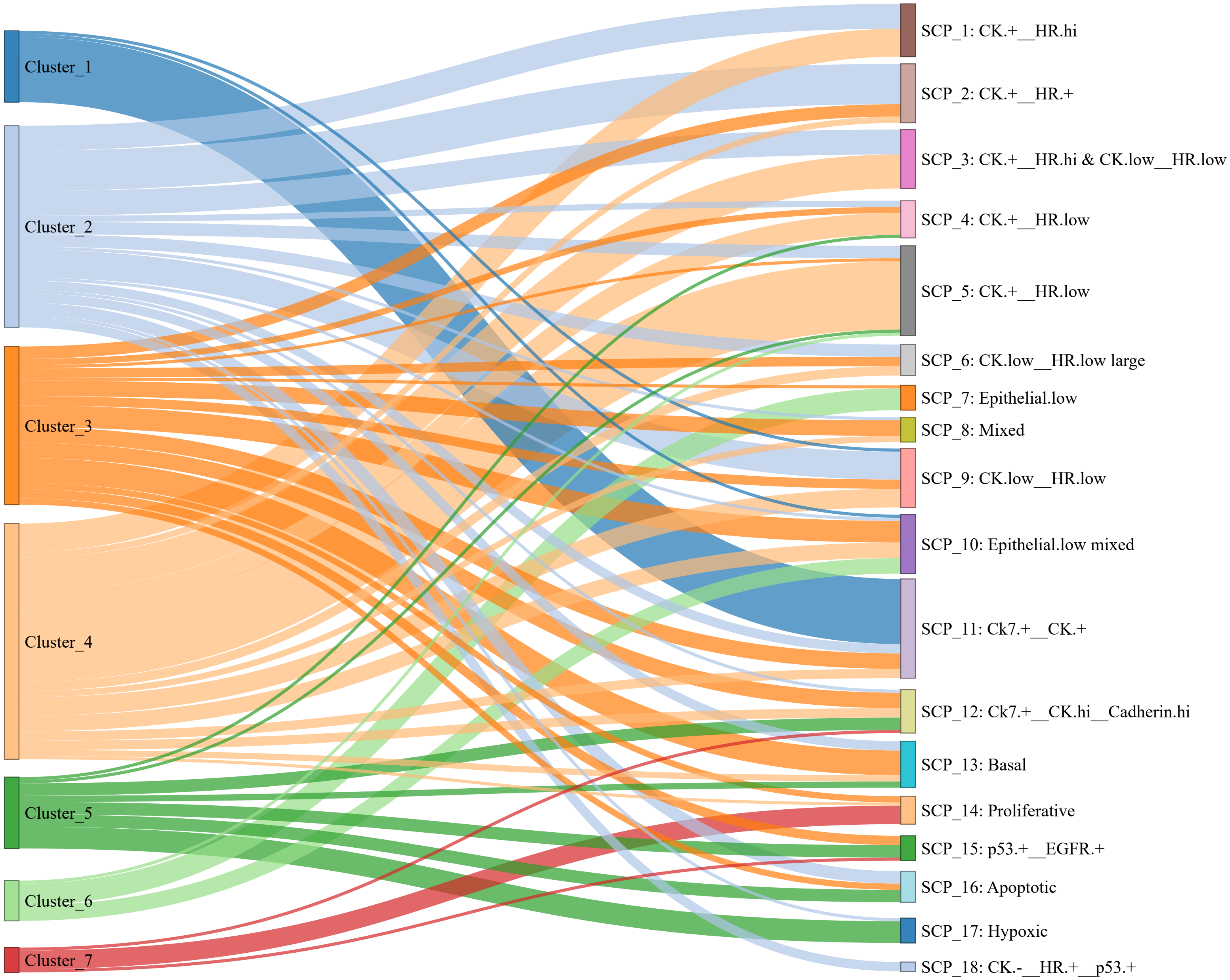

### Supplemental Figure 2

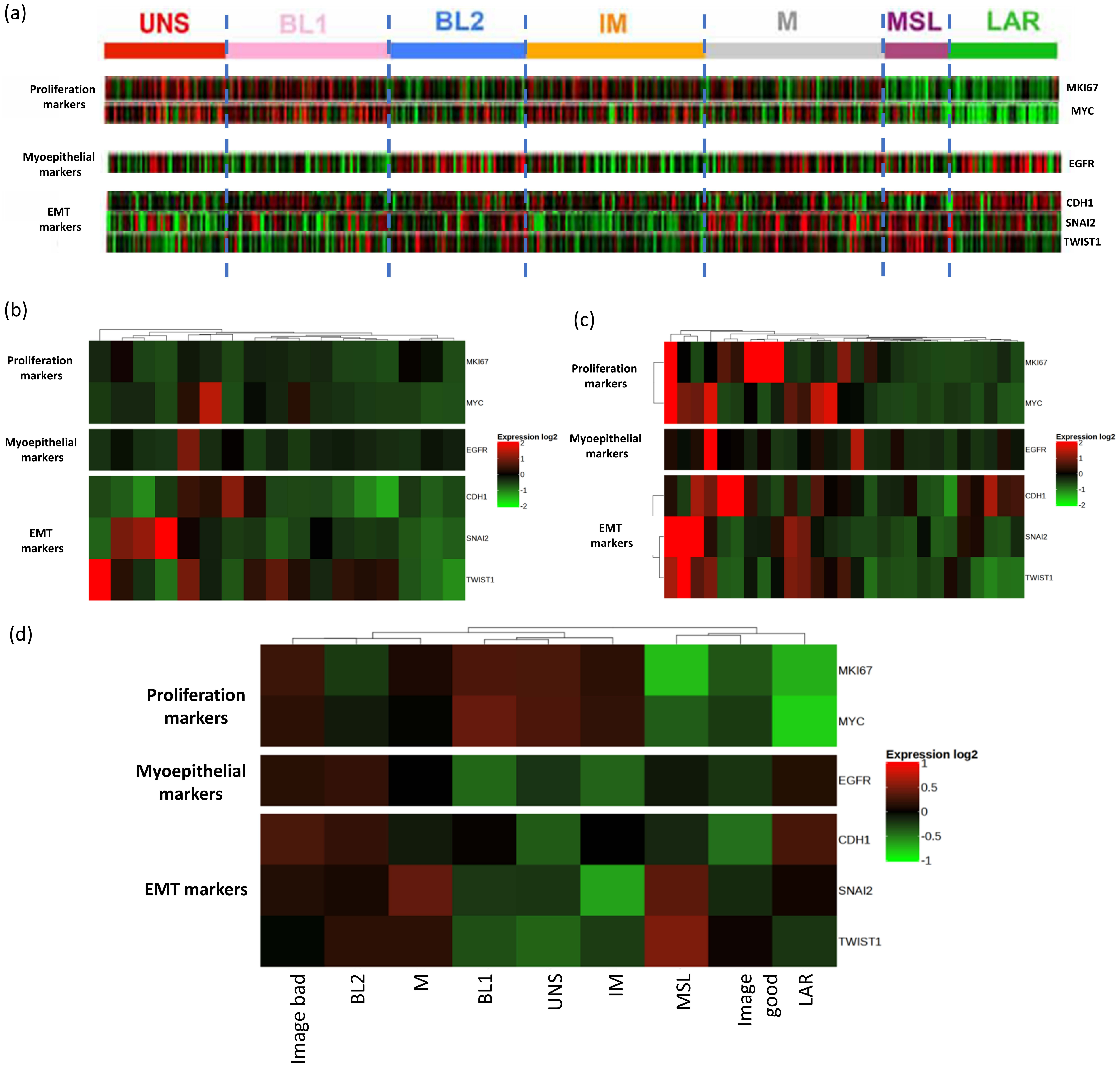

### Supplemental Figure 3

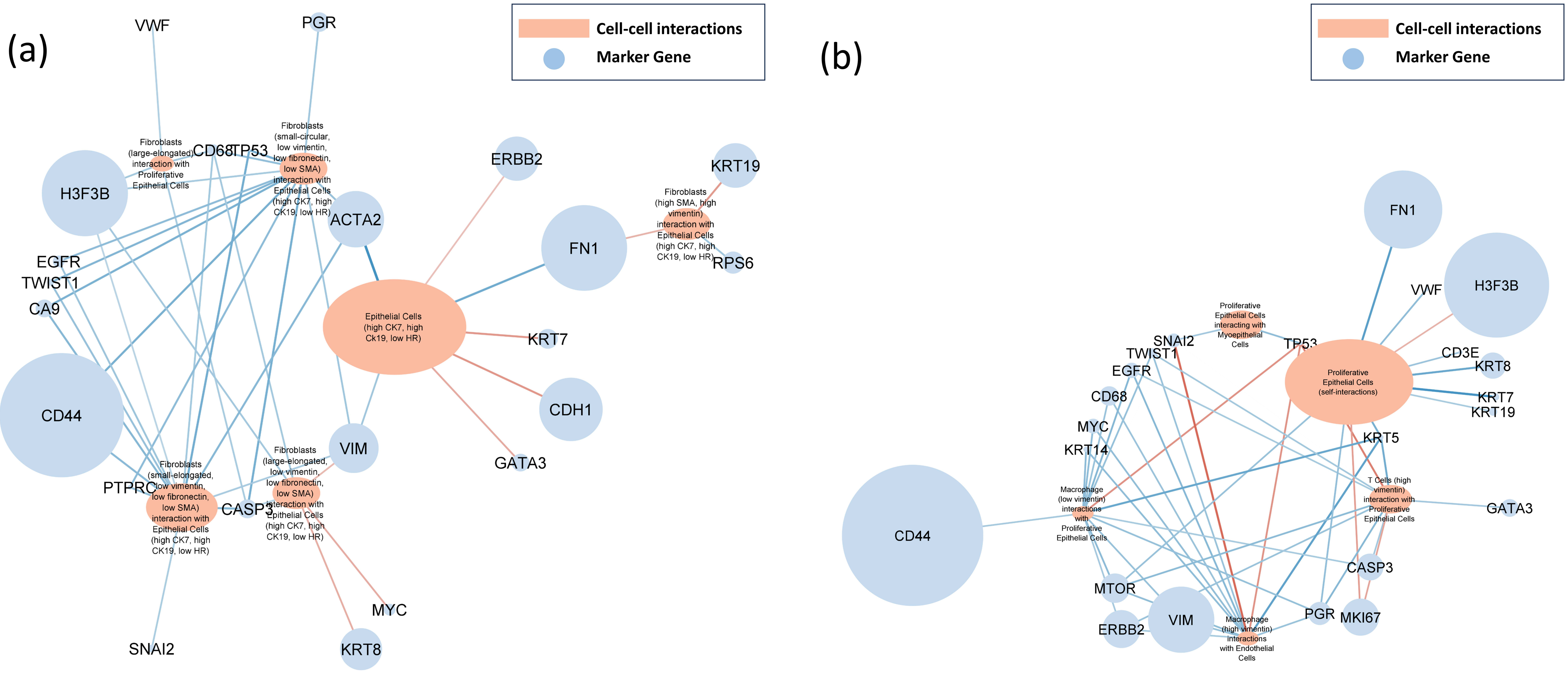
