## Supplemental Table 1 for "Deep-learning and transfer learning identify new breast cancer survival subtypes from single-cell imaging data"

Table S1

| Clinical Features | | | | | | | | | | Subtypes | | | |
| --- | --- | --- | --- | --- | --- | --- | --- | --- | --- | --- | --- | --- | --- |
| | Tumor Grade | | | ER Status | | PR Status | | HER2 Status | | Luminal A | Luminal B | TNBC | HER2 Enriched |
| **Total** | **I** | **II** | **III** | + | **-** | + | **-** | + | **-** | | | | |
| **259** | **34** | **109** | **116** | **191** | **68** | **143** | **116** | **49** | **210** | **166** | **26** | **44** | **23** |
