## Supplemental Table 2 for "Deep-learning and transfer learning identify new breast cancer survival subtypes from single-cell imaging data"

Table S2

| # | Cell Phenotype | Description |
|---|---|---|
| **1** | B Cell | Immune cells showing expression of CD20 (B-Cell biomarker), CD45 (pan-immune biomarker), and Vimentin Expression. |
| **2** | T and B Cell | Immune cell cluster (T Cells appearing together with B cells). Hierarchical Clustering is not able to unmix these cells based on the biomarkers. These cells express CD3 (T-Cell biomarker), CD45 (pan-immune biomarker), and CD20 (B-Cell biomarker). This cluster has a low Vimentin expression. |
| **3** | T $Cell_1$ | Immune cells showing expression of CD3(T Cell Biomarker) and CD45 (pan-immune biomarker). This cluster has a low Vimentin expression. |
| **4** | $Macrophage_1$ | Immune cells expressing CD68 (macrophage biomarker). This cluster has a high level of Vimentin expression. |
| **5** | T $Cell_2$ | Immune cells showing expression of CD3(T Cell Biomarker) and CD45 (pan-immune biomarker). These T Cells have high levels of vimentin expression as compared to the T Cells in Cluster 3. |
| **6** | $Macrophage_2$ | Immune cells expressing CD68 (macrophage biomarker). These macrophages have low levels of vimentin expression compared to $Macrophage_1$ |
| **7** | Endothelial cell | These cells form the barrier between blood vessels and tissues. Expressing vWF(von Willebrand factor - endothelial marker), CD31 (endothelial marker). These cells also show high Vimentin Expression. |
| **8** | $Vimentin^{hi}$ Fibroblast | Fibroblasts cells with high vimentin expression, low smooth muscle actin (SMA) expression, low fibronectin expression. These cells also express high cMYC (a proto-oncogene). |
| **9** | Small circular Fibroblast | These fibroblasts cells are small and circular and show low expression of Vimentin, Fibronectin, and SMA. |
| **10** | Small elongated Fibroblast | These are fibroblasts cells small in size and elongated (high eccentricity) and show low expression of Vimentin, Fibronectin, and SMA. |
| **11** | $Fibronectin^{hi}$ Fibroblast | Fibroblasts cells with high fibronectin expression, low smooth muscle actin (SMA) expression, low vimentin expression. |
| **12** | Large Elongated Fibroblast | These fibroblasts cells are larger and elongated and show low Vimentin, Fibronectin, and SMA expression. |
| **13** | $SMA^{hi}$ $Vimentin^{hi}$ Fibroblast | Fibroblasts cells with high SMA expression, high vimentin expression, and low fibronectin expression. They also show an increased expression of CD68. |
| **14** | Hypoxic epithelial | These epithelial cells have a high expression of CAIX (Carbonic Anhydrase), which is a hypoxia marker. Rest other epithelial markers such as cytokeratins, e-cadherins, ER, PR have a deficient expression. |
| **15** | Apoptotic epithelial | These epithelial cells have a high expression of PARP (Poly (ADP-ribose) polymerase) and Caspase 3, markers for apoptosis. These cells also show an increased expression of p53 and EGFR (epidermal growth factor receptor). Rest other epithelial markers such as cytokeratin, e-cadherins, ER, PR have a deficient expression. |
| **16** | Proliferative epithelial | These epithelial cells have high expressions of Ki67 (nuclear protein associated with cellular proliferation), pHH3 (Phosphorylated Histone H3), and Phospho-S6 |

| | | |
|---|---|---|
| | | ribosomal protein. Rest other epithelial markers such as cytokeratins, e-cadherins, ER, PR have a deficient expression. |
| **17** | p53 $^{+}$ EGFR $^{+}$ epithelial | These epithelial cells have high expressions of p53 and EGFR (epidermal growth factor receptor). Other significantly expressed biomarkers are CAIX (Carbonic Anhydrase), cMYC (a proto-oncogene). Rest other epithelial markers such as cytokeratins, e-cadherins, ER, PR have a deficient expression. |
| **18** | Basal CK epithelial | These epithelial cells have high expressions of CK5 and CK14 (basal cytokeratin biomarkers). Rest other epithelial markers such as e-cadherins, ER, PR have an extremely low expression. |
| **19** | CK7 $^{+}$ CK $^{hi}$ Cadherin $^{hi}$ epithelial | These epithelial cells have high expressions of CK7, other luminal cytokeratins (CK8/18, CK19 biomarkers), and E/P Cadherins. Other epithelial markers, ER, PR have an extremely low expression. |
| **20** | CK7 $^{+}$CK $^{+}$ epithelial | These epithelial cells have high expressions of CK19, other luminal cytokeratins (CK7 biomarkers). Rest other epithelial markers such as e-cadherins, ER, PR have an extremely low expression. |
| **21** | Epithelial $^{low}$ | These epithelial cells have almost negligible expressions of any epithelial markers such as cytokeratins, e-cadherins, ER, PR. Also, they do not express other markers such as p53, EGFR, CAIX, Ki67, PHH3. |
| **22** | CK $^{low}$ HR $^{low}$ epithelial | These epithelial cells have low expressions of epithelial markers such as cytokeratins, ER, PR. Also, they do not express other markers such as p53, EGFR, CAIX, Ki67, PHH3. |
| **23** | CK$^{+}$ HR $^{hi}$ epithelial | These epithelial cells have very high expressions of epithelial markers such as luminal cytokeratins (CK 8/18, CK 19), pan-cytokeratin, ER, PR. They also express high HER2. However, luminal CK 7 is absent in these cells. |
| **24** | CK$^{+}$HR$^{+}$ epithelial | These epithelial cells have very high expressions of epithelial markers such as luminal cytokeratins (CK 8/18, CK 19), pan-cytokeratin. They do not have an increased expression of ER, PR. They also express high HER2. However, luminal CK 7 is absent in these cells. |
| **25** | CK$^{+}$HR $^{low}$ epithelial | These epithelial cells have very high expressions of epithelial markers such as luminal cytokeratins (CK 8/18, CK 19), pan-cytokeratin. However, they have extremely low ER, PR, HER2 expressions. Luminal CK 7 is absent in these cells. |
| **26** | CK $^{low}$ HR$^{hi}$ p53$^{+}$ epithelial | These epithelial cells have very low expressions of epithelial markers such as luminal cytokeratins (CK 8/18, CK 19), pan-cytokeratin. However, they have high ER, PR, HER2 expression. They also express a significant amount of p53, EGFR, PARP, and Caspase 3 |
| **27** | Myoepithelial | These epithelial cells form semi-continuous protective sheets separating the human breast epithelium and the surrounding stroma and show higher expression of SMA, Luminal Cytokeratins, Basal Cytokeratins, pan Cytokeratins biomarkers. |
