## Supplemental Table 3 for "Deep-learning and transfer learning identify new breast cancer survival subtypes from single-cell imaging data"

Table S3

| Rank | Cellular Phenotype | Importance Score | Type |
|---|---|---|---|
| 1 | $Macrophage_2$ | 1.000 | Immune |
| 2 | T and B Cell | 0.692 | Immune |
| 3 | $CK^+HR^+$ | 0.631 | Epithelial |
| 4 | $CK7^+CK^+$ | 0.576 | Epithelial |
| 5 | Endothelial | 0.546 | Stromal |
| 6 | Hypoxic | 0.543 | Epithelial |
| 7 | $CK^+HR^{low}$ | 0.508 | Epithelial |
| 8 | $CK^+HR^{hi}$ | 0.507 | Epithelial |
