## Supplemental Table 4 for "Deep-learning and transfer learning identify new breast cancer survival subtypes from single-cell imaging data"

Table S4

| Rank | Cell Phenotype 1 | Cell Phenotype 2 | Importance Score | Interaction Type |
|---|---|---|---|---|
| 1 | T and B Cell | $CK^{low}HR^{low}$ | 1.000 | Immune-Epithelial |
| 2 | Small Elongated Fibroblast | $CK7^{+}CK^{+}$ | 0.966 | Stromal-Epithelial |
| 3 | $Macrophage_1$ | $CK^{low}HR^{hi}p53^{+}$ | 0.965 | Immune-Epithelial |
| 4 | $Vimentin^{hi}$ Fibroblast | Small Circular Fibroblast | 0.964 | Stromal-Stromal |
| 5 | Large Elongated Fibroblast | Proliferative | 0.922 | Stromal-Epithelial |
| 6 | $SMA^{hi}Vimentin^{hi}$ Fibroblast | $CK7^{+}CK^{+}$ | 0.892 | Stromal-Epithelial |
| 7 | B Cell | $Epithelial^{low}$ | 0.885 | Immune-Epithelial |
| 8 | T and B Cell | Hypoxic | 0.867 | Immune-Epithelial |
| 9 | $Macrophage_2$ | Small Circular Fibroblast | 0.853 | Immune-Stromal |
| 10 | $Macrophage_2$ | Small Elongated Fibroblast | 0.848 | Immune-Stromal |
| 11 | $SMA^{hi}Vimentin^{hi}$ Fibroblast | $CK^{low}HR^{hi}p53^{+}$ | 0.846 | Stromal-Epithelial |
| 12 | Small Circular Fibroblast | $CK7^{+}CK^{+}$ | 0.841 | Stromal-Epithelial |
| 13 | $Vimentin^{hi}$ Fibroblast | $CK^{low}HR^{hi}p53^{+}$ | 0.823 | Stromal-Epithelial |
| 14 | T and B Cell | $CK^{low}HR^{hi}p53^{+}$ | 0.819 | Immune-Epithelial |
| 15 | $Macrophage_1$ | Endothelial | 0.814 | Immune-Stromal |
| 16 | T and B Cell | Endothelial | 0.809 | Immune-Stromal |
| 17 | Endothelial | $CK^{low}HR^{hi}p53^{+}$ | 0.808 | Stromal-Epithelial |
| 18 | $SMA^{hi}Vimentin^{hi}$ | $Epithelial^{low}$ | 0.795 | Stromal-Epithelial |
| 19 | T $Cell_2$ | Proliferative | 0.792 | Immune-Epithelial |
| 20 | $Macrophage_1$ | $CK^{+}HR^{hi}$ | 0.783 | Immune-Epithelial |
| 21 | Small Circular Fibroblast | $CK^{+}HR^{hi}$ | 0.773 | Stromal-Epithelial |
| 22 | Small Elongated Fibroblast | $CK^{+}HR^{+}$ | 0.769 | Stromal-Epithelial |
| 23 | B Cell | $Fibronectin^{hi}$ Fibroblast | 0.766 | Immune-Stromal |
| 24 | $Vimentin^{hi}$ Fibroblast | $CK^{+}HR^{+}$ | 0.765 | Stromal-Epithelial |
| 25 | $SMA^{hi}Vimentin^{hi}$ | $CK^{low}HR^{low}$ | 0.765 | Stromal-Epithelial |
| 26 | $SMA^{hi}Vimentin^{hi}$ | Apoptotic | 0.764 | Stromal-Epithelial |
| 27 | Small Circular Fibroblast | $CK7^{+}CK^{hi}cadherin^{hi}$ | 0.760 | Stromal-Epithelial |
| 28 | T $Cell_2$ | Apoptotic | 0.760 | Immune-Epithelial |
| 29 | Large Elongated Fibroblast | $CK7^{+}CK^{+}$ | 0.751 | Immune-Stromal |
| 30 | $Macrophage_2$ | Proliferative | 0.751 | Immune-Epithelial |
