## Supplemental Table 5 for "Deep-learning and transfer learning identify new breast cancer survival subtypes from single-cell imaging data"

Table S5

| Rank | Cell Phenotype 1 | Cell Phenotype 2 | Importance Score | Interaction Type |
|---|---|---|---|---|
| 1 | Proliferative | Proliferative | 1.000 | Epithelial-Epithelial |
| 2 | $Epithelial^{low}$ | $CK^{low}$ $HR^{hi}$ $p53^{+}$ | 0.870 | |
| 3 | $CK^{+}$ $HR^{+}$ | $CK^{+}$ $HR^{low}$ | 0.811 | |
| 4 | Hypoxic | $CK^{+}$ $HR^{+}$ | 0.806 | |
| 5 | Basal CK | $CK7^{+}$ $CK^{hi}$ $cadherin^{hi}$ | 0.789 | |
| 6 | $CK^{low}$ $HR^{low}$ | $CK^{+}$ $HR^{+}$ | 0.788 | |
| 7 | Proliferative | Basal CK | 0.783 | |
| 8 | Hypoxic | $Epithelium^{low}$ | 0.775 | |
| 9 | Proliferative | $CK^{+}$ $HR^{low}$ | 0.769 | |
| 10 | Proliferative | Myoepithelial | 0.766 | |
| 11 | $Epithelial^{low}$ | $Epithelial^{low}$ | 0.766 | |
| 12 | $CK^{+}$ $HR^{+}$ | Myoepithelial | 0.763 | |
