## Supplemental Table 6 for "Deep-learning and transfer learning identify new breast cancer survival subtypes from single-cell imaging data"

Table S6

| Distribution of Patient in each Cluster Clinical Features | | | | | | | | | | | | | | |
|---|---|---|---|---|---|---|---|---|---|---|---|---|---|---|
| Clinical Features | | | | | | | | | | | Subtypes | | | |
| **Cluster ID** | **No. Of Patients** | **Tumor Grade** | | | **ER Status** | | **PR Status** | | **HER2 Status** | | | | | |
| | | **I** | **II** | **III** | + | - | + | - | + | - | **Luminal A** | **Luminal B** | **TNBC** | **HER2 Enriched** |
| **1** | 23 | 6 | 9 | 8 | 14 | 9 | 12 | 11 | 7 | 16 | 14 | 0 | 2 | 7 |
| **2** | 65 | 7 | 35 | 23 | 59 | 6 | 45 | 20 | 13 | 52 | 48 | 11 | 4 | 2 |
| **3** | 51 | 4 | 15 | 32 | 27 | 24 | 19 | 32 | 12 | 39 | 21 | 6 | 18 | 6 |
| **4** | 76 | 16 | 41 | 19 | 71 | 5 | 55 | 21 | 8 | 68 | 65 | 7 | 3 | 1 |
| **5** | 23 | 0 | 3 | 20 | 9 | 14 | 5 | 18 | 6 | 17 | 8 | 1 | 9 | 5 |
| 6 | 13 | 0 | 6 | 7 | 9 | 4 | 5 | 8 | 3 | 10 | 8 | 1 | 2 | 2 |
| **7** | 8 | 1 | 0 | 7 | 2 | 6 | 2 | 6 | 0 | 8 | 2 | 0 | 6 | 0 |
