## Supplemental Table 7 for "Deep-learning and transfer learning identify new breast cancer survival subtypes from single-cell imaging data"

| Gene Symbol | logFC | AveExpr | t | P.Value | adj.P.Val |
|---|---|---|---|---|---|
| CA9 | -1.66532 | 1.181947 | -3.43789 | 0.001317 | 0.037967 |
| GATA3 | 1.735083 | 1.894308 | 3.208068 | 0.002531 | 0.037967 |
| PTPRC | 2.099658 | 2.148328 | 3.016766 | 0.004288 | 0.042875 |
| CDH1 | -1.29972 | 2.695109 | -2.6223 | 0.012046 | 0.090348 |
| MKI67 | -0.39527 | 0.455639 | -1.98155 | 0.053973 | 0.323837 |
| PGR | 0.31247 | 0.670969 | 1.870176 | 0.068305 | 0.341527 |
| CD68 | 0.263879 | 0.694707 | 1.555042 | 0.127297 | 0.545558 |
| H3F3B | -2.03734 | 8.670026 | -1.48123 | 0.145868 | 0.547003 |
| MYC | -0.25257 | 0.561824 | -1.32181 | 0.193251 | 0.604844 |
| MTOR | 0.32937 | 0.957116 | 1.276558 | 0.208638 | 0.604844 |
| KRT14 | 0.127985 | 0.24281 | 1.239867 | 0.221776 | 0.604844 |
| EGFR | -0.29323 | 0.633435 | -1.14713 | 0.257698 | 0.644245 |
| ESR1 | -0.04331 | 0.154521 | -0.99907 | 0.323374 | 0.728471 |
| ERBB2 | -0.20024 | 1.177114 | -0.96502 | 0.339953 | 0.728471 |
| SNAI2 | -0.07591 | 0.292052 | -0.87052 | 0.388872 | 0.766365 |
| CASP3 | 0.122374 | 0.87393 | 0.799978 | 0.42814 | 0.766365 |
| KRT7 | -0.05051 | 0.187082 | -0.77142 | 0.444694 | 0.766365 |
| CD3E | -0.02687 | 0.119141 | -0.74587 | 0.459819 | 0.766365 |
| ACTA2 | -0.07116 | 0.331779 | -0.60659 | 0.547323 | 0.864193 |
| TP53 | 0.007247 | 0.093337 | 0.395215 | 0.694646 | 0.94636 |
| VIM | 0.35772 | 4.605131 | 0.311459 | 0.756963 | 0.94636 |
| KRT5 | 0.016114 | 0.184095 | 0.288969 | 0.774 | 0.94636 |
| VWF | -0.00535 | 0.109268 | -0.28682 | 0.775636 | 0.94636 |
| CD44 | -1.57545 | 33.46401 | -0.25769 | 0.79788 | 0.94636 |
| MS4A1 | -0.01068 | 0.142771 | -0.23376 | 0.816284 | 0.94636 |
| KRT19 | -0.01825 | 0.381231 | -0.17763 | 0.85985 | 0.94636 |
| TWIST1 | 0.006286 | 0.238186 | 0.1507 | 0.88092 | 0.94636 |
| KRT8 | 0.048855 | 0.645513 | 0.147705 | 0.883269 | 0.94636 |
| FN1 | 0.009486 | 0.345973 | 0.072506 | 0.942537 | 0.975038 |
| RPS6 | -0.00023 | 0.9968 | -0.00104 | 0.999173 | 0.999173 |
