## Supplemental Table 8 for "Deep-learning and transfer learning identify new breast cancer survival subtypes from single-cell imaging data"

| Gene Symbol | logFC | AveExpr | t | P.Value | adj.P.Val |
|---|---|---|---|---|---|
| KRT7 | 0.621095 | 0.252991 | 3.392664 | 0.000867 | 0.022652 |
| CASP3 | -0.30149 | 0.716597 | -3.18383 | 0.001738 | 0.022652 |
| ERBB2 | -0.71272 | 1.428197 | -3.10159 | 0.002265 | 0.022652 |
| MTOR | -0.47299 | 1.085437 | -2.92243 | 0.003961 | 0.025079 |
| H3F3B | -3.88308 | 8.254407 | -2.90479 | 0.00418 | 0.025079 |
| TP53 | -0.02037 | 0.068909 | -2.74515 | 0.00672 | 0.02899 |
| GATA3 | -0.43225 | 0.831243 | -2.7429 | 0.006764 | 0.02899 |
| ACTA2 | 1.450311 | 2.908928 | 2.64274 | 0.009017 | 0.030062 |
| FN1 | 0.966874 | 1.939285 | 2.642688 | 0.009019 | 0.030062 |
| CDH1 | -1.49285 | 3.209465 | -2.53702 | 0.012109 | 0.036327 |
| ESR1 | -0.20296 | 0.333834 | -2.36442 | 0.019223 | 0.052427 |
| TWIST1 | -0.06338 | 0.228428 | -2.14722 | 0.033239 | 0.082595 |
| VWF | -0.02091 | 0.091888 | -2.09964 | 0.037285 | 0.082595 |
| KRT19 | 0.611251 | 1.045851 | 2.085735 | 0.038544 | 0.082595 |
| SNAI2 | -0.10615 | 0.223992 | -1.89526 | 0.05981 | 0.11962 |
| CD68 | -0.20877 | 0.480148 | -1.85104 | 0.065955 | 0.123665 |
| KRT14 | -0.05732 | 0.15019 | -1.72856 | 0.085763 | 0.143635 |
| RPS6 | -0.27315 | 0.831391 | -1.70041 | 0.090942 | 0.143635 |
| PTPRC | -0.08485 | 0.20329 | -1.70026 | 0.090969 | 0.143635 |
| PGR | -0.4174 | 0.99885 | -1.63683 | 0.103576 | 0.155364 |
| CA9 | 0.215853 | 0.747 | 1.449928 | 0.14898 | 0.212829 |
| CD3E | -0.01952 | 0.075158 | -1.39362 | 0.165311 | 0.225424 |
| KRT5 | -0.01767 | 0.131671 | -1.00806 | 0.314905 | 0.410746 |
| KRT8 | 0.487363 | 1.86885 | 0.962083 | 0.337418 | 0.421773 |
| VIM | -0.45621 | 2.978811 | -0.89505 | 0.372069 | 0.446483 |
| CD44 | -5.46826 | 22.58901 | -0.74825 | 0.455379 | 0.525438 |
| MS4A1 | -0.05775 | 0.11631 | -0.67831 | 0.498525 | 0.553917 |
| MYC | -0.01087 | 0.407169 | -0.13294 | 0.894403 | 0.926926 |
| EGFR | -0.00688 | 0.370803 | -0.13088 | 0.896028 | 0.926926 |
| MKI67 | 0.002209 | 0.138045 | 0.056161 | 0.955281 | 0.955281 |
