## Supplemental Table 9 for "Deep-learning and transfer learning identify new breast cancer survival subtypes from single-cell imaging data"

| Gene Name | logFC | AveExpr | t | P.Value | adj.P.Val |
|---|---|---|---|---|---|
| KRT5 | 0.359085 | 0.353412 | 3.602914 | 0.000415 | 0.008454 |
| CA9 | -0.733360 | -0.780730 | -3.496940 | 0.000604 | 0.008454 |
| GATA3 | 0.944708 | -0.016850 | 3.098935 | 0.002282 | 0.021295 |
| CD3E | -0.451420 | 0.369958 | -2.201850 | 0.029055 | 0.203382 |
| MS4A1 | -0.445460 | 0.301218 | -2.030820 | 0.043868 | 0.245663 |
| KRT8 | -0.261330 | 0.585154 | -1.725830 | 0.086238 | 0.375687 |
| EGFR | 0.283720 | 0.989292 | 1.684717 | 0.093922 | 0.375687 |
| TP53 | -0.374510 | 0.199245 | -1.616030 | 0.107987 | 0.377956 |
| PTPRC | -0.335660 | 0.323452 | -1.474140 | 0.142338 | 0.423541 |
| MYC | 0.293505 | 0.615234 | 1.441721 | 0.151265 | 0.423541 |
| ACTA2 | 0.316745 | -0.131780 | 1.322205 | 0.187919 | 0.478338 |
| H3F3B | 0.183001 | 0.058156 | 1.009132 | 0.314380 | 0.717750 |
| CD44 | 0.181941 | 0.170509 | 0.920988 | 0.358393 | 0.717750 |
| CDH1 | 0.144242 | -0.144540 | 0.845772 | 0.398897 | 0.717750 |
| CD68 | 0.163085 | 0.233515 | 0.754943 | 0.451353 | 0.717750 |
| FN1 | 0.151448 | -0.245670 | 0.744126 | 0.457852 | 0.717750 |
| VIM | -0.145540 | 0.583144 | -0.711480 | 0.477788 | 0.717750 |
| MKI67 | 0.120782 | 0.761689 | 0.687042 | 0.493015 | 0.717750 |
| PGR | 0.176797 | -2.415230 | 0.672580 | 0.502150 | 0.717750 |
| ERBB2 | -0.085600 | -0.730870 | -0.656080 | 0.512678 | 0.717750 |
| CASP3 | -0.116840 | -1.556450 | -0.584980 | 0.559356 | 0.745808 |
| VWF | 0.114310 | -0.418070 | 0.485462 | 0.627989 | 0.799259 |
| RPS6 | -0.090500 | 0.383519 | -0.351920 | 0.725345 | 0.864719 |
| ESR1 | -0.041970 | -1.605010 | -0.302680 | 0.762516 | 0.864719 |
| KRT7 | -0.057680 | 0.814242 | -0.276940 | 0.782170 | 0.864719 |
| KRT19 | 0.064105 | -0.544870 | 0.249926 | 0.802953 | 0.864719 |
| KRT14 | 0.038968 | 0.403564 | 0.191417 | 0.848433 | 0.867726 |
| SNAI2 | -0.042370 | 0.135287 | -0.166810 | 0.867726 | 0.867726 |
