## Supplemental Table 10 for "Deep-learning and transfer learning identify new breast cancer survival subtypes from single-cell imaging data"

| Gene Name | logFC | AveExpr | t | P.Value | adj.P.Val |
|---|---|---|---|---|---|
| KRT14 | 0.866476 | -0.059990 | 7.098703 | 3.48E-12 | 9.74E-11 |
| SNAI2 | 0.783525 | -0.030140 | 6.636017 | 7.07E-11 | 9.90E-10 |
| KRT7 | 0.787224 | -0.103080 | 6.549817 | 1.22E-10 | 1.14E-09 |
| ACTA2 | 0.678161 | 0.027382 | 5.873735 | 6.97E-09 | 4.88E-08 |
| VWF | 0.646944 | 0.152885 | 5.629324 | 2.75E-08 | 1.49E-07 |
| VIM | 0.653774 | -0.093400 | 5.602645 | 3.18E-08 | 1.49E-07 |
| MYC | 0.467220 | -0.058380 | 4.031164 | 0.000063 | 0.000250 |
| CDH1 | -0.506140 | -0.049480 | -3.872330 | 0.000119 | 0.000418 |
| EGFR | 0.403637 | -0.249010 | 3.765384 | 0.000182 | 0.000567 |
| PTPRC | 0.405830 | -0.031410 | 3.471977 | 0.000553 | 0.001548 |
| MKI67 | -0.411220 | -0.271530 | -3.399980 | 0.000717 | 0.001826 |
| ESR1 | -0.225690 | 0.446677 | -3.262840 | 0.001164 | 0.002716 |
| KRT5 | 0.395878 | -0.181310 | 3.227836 | 0.001314 | 0.002829 |
| KRT8 | -0.380290 | 0.114790 | -3.202750 | 0.001431 | 0.002863 |
| GATA3 | -0.240510 | 0.378847 | -3.093340 | 0.002069 | 0.003862 |
| CD3E | 0.350318 | -0.074010 | 2.877213 | 0.004151 | 0.007264 |
| MS4A1 | 0.326671 | -0.136920 | 2.846465 | 0.004568 | 0.007524 |
| KRT19 | -0.288910 | 0.098399 | -2.590640 | 0.009806 | 0.015254 |
| PGR | 0.261333 | 0.377742 | 2.499440 | 0.012698 | 0.018713 |
| RPS6 | 0.252696 | -0.064950 | 2.259145 | 0.024223 | 0.033912 |
| ERBB2 | 0.149491 | -0.186800 | 2.104345 | 0.035753 | 0.047272 |
| CASP3 | -0.229720 | -0.187110 | -2.088730 | 0.037142 | 0.047272 |
| CA9 | -0.184530 | -0.477180 | -1.784310 | 0.074865 | 0.091140 |
| H3F3B | -0.209870 | -0.018020 | -1.728320 | 0.084432 | 0.098504 |
| FN1 | 0.168311 | -0.020360 | 1.360655 | 0.174120 | 0.195015 |
| CD44 | 0.114186 | 0.031027 | 0.937303 | 0.348970 | 0.375814 |
| TP53 | 0.078552 | 0.078036 | 0.813627 | 0.416173 | 0.431587 |
| CD68 | 0.043611 | -0.054470 | 0.354237 | 0.723282 | 0.723282 |
