## Supplemental Table 11 for "Deep-learning and transfer learning identify new breast cancer survival subtypes from single-cell imaging data"

| Gene Name | logFC | AveExpr | t | P.Value | adj.P.Val |
|---|---|---|---|---|---|
| ACTA2 | 0.573868 | 0.077270 | 3.670104 | 2.87E-04 | 7.61E-03 |
| KRT7 | -0.549600 | 0.484024 | -3.515140 | 5.07E-04 | 7.61E-03 |
| GATA3 | 0.638064 | -0.102740 | 3.319498 | 1.01E-03 | 1.01E-02 |
| H3F3B | 0.278453 | -0.369990 | 1.992902 | 4.72E-02 | 2.94E-01 |
| EGFR | -0.338870 | 1.203013 | -1.976670 | 4.90E-02 | 2.94E-01 |
| MS4A1 | 0.290526 | 0.117949 | 1.779548 | 7.62E-02 | 3.81E-01 |
| CD68 | 0.266183 | 0.410858 | 1.592282 | 0.112372 | 0.481593 |
| TP53 | 0.158366 | -1.685070 | 1.428578 | 0.154163 | 0.578111 |
| VWF | 0.190017 | -0.222290 | 1.277852 | 0.202287 | 0.612230 |
| FN1 | 0.185938 | 0.086585 | 1.272786 | 0.204077 | 0.612230 |
| KRT19 | -0.227440 | -0.582490 | -1.178710 | 0.239444 | 0.653029 |
| KRT8 | -0.204340 | -0.898690 | -1.099790 | 0.272301 | 0.680753 |
| CDH1 | -0.125990 | -0.211900 | -0.945570 | 0.345129 | 0.751691 |
| RPS6 | -0.135820 | 0.079517 | -0.907200 | 0.365026 | 0.751691 |
| ERBB2 | 0.109310 | -0.729280 | 0.886892 | 0.375845 | 0.751691 |
| PTPRC | 0.146653 | 0.539348 | 0.839543 | 0.401832 | 0.753434 |
| CA9 | -0.156040 | 0.910100 | -0.764220 | 0.445338 | 0.776903 |
| KRT5 | -0.125250 | 0.950768 | -0.697810 | 0.485832 | 0.776903 |
| MTOR | -0.098440 | 0.138333 | -0.641760 | 0.521519 | 0.776903 |
| MYC | -0.088930 | 0.513788 | -0.596690 | 0.551162 | 0.776903 |
| CD44 | 0.074794 | 0.084156 | 0.569977 | 0.569119 | 0.776903 |
| MKI67 | 0.091584 | 0.836039 | 0.539762 | 0.589761 | 0.776903 |
| ESR1 | 0.026827 | -1.662260 | 0.531267 | 0.595626 | 0.776903 |
| VIM | 0.040330 | 0.477915 | 0.314419 | 0.753421 | 0.909424 |
| SNAI2 | 0.047458 | 0.241982 | 0.271032 | 0.786552 | 0.909424 |
| CASP3 | -0.038570 | 0.454359 | -0.268930 | 0.788168 | 0.909424 |
| CD3E | -0.031100 | 0.466719 | -0.180230 | 0.857091 | 0.910779 |
| KRT14 | -0.025880 | 0.362967 | -0.151640 | 0.879570 | 0.910779 |
| PGR | -0.00352 | -0.91613 | -0.12531 | 0.900364 | 0.910779 |
| TWIST1 | -0.01595 | 0.154403 | -0.11215 | 0.910779 | 0.910779 |
