## Supplemental Table 12 for "Deep-learning and transfer learning identify new breast cancer survival subtypes from single-cell imaging data"

| Gene Name | logFC | AveExpr | t | P.Value | adj.P.Val |
|---|---|---|---|---|---|
| KRT7 | 0.977340 | -0.097440 | 3.803803 | 1.49E-04 | 4.46E-03 |
| ACTA2 | 0.703218 | -0.001900 | 3.603348 | 3.25E-04 | 4.88E-03 |
| CDH1 | -0.674130 | -0.543030 | -3.399420 | 6.95E-04 | 6.95E-03 |
| VIM | 0.845528 | -0.193060 | 3.084499 | 2.08E-03 | 1.56E-02 |
| MS4A1 | -0.592870 | -0.177460 | -2.577770 | 1.00E-02 | 6.03E-02 |
| VWF | 0.191635 | 0.053709 | 1.934711 | 5.32E-02 | 2.66E-01 |
| GATA3 | 0.096992 | 0.455363 | 1.832000 | 0.067168 | 0.287864 |
| KRT14 | 0.154100 | -0.197550 | 1.588594 | 0.112383 | 0.374411 |
| SNAI2 | 0.148167 | -0.055810 | 1.562958 | 0.118293 | 0.374411 |
| TP53 | -0.129190 | 0.007260 | -1.488630 | 0.136816 | 0.374411 |
| CA9 | -0.096260 | -0.256750 | -1.486850 | 0.137284 | 0.374411 |
| CD3E | -0.119040 | -0.157230 | -1.363400 | 0.172979 | 0.401474 |
| KRT19 | 0.118648 | 0.097706 | 1.326799 | 0.184796 | 0.401474 |
| KRT5 | 0.106327 | -0.181740 | 1.283057 | 0.199689 | 0.401474 |
| KRT8 | 0.104671 | 0.131979 | 1.280070 | 0.200737 | 0.401474 |
| FN1 | 0.090496 | -0.083140 | 0.950817 | 0.341865 | 0.622803 |
| CD68 | 0.083207 | -0.109750 | 0.891749 | 0.372684 | 0.622803 |
| ERBB2 | 0.051047 | -0.191510 | 0.889889 | 0.373682 | 0.622803 |
| RPS6 | 0.081818 | 0.021178 | 0.837080 | 0.402693 | 0.635832 |
| PGR | 0.070527 | 0.308679 | 0.720813 | 0.471147 | 0.688093 |
| EGFR | -0.046280 | -0.306160 | -0.691850 | 0.489147 | 0.688093 |
| CD44 | 0.066715 | 0.038655 | 0.667443 | 0.504602 | 0.688093 |
| H3F3B | -0.049260 | 0.133718 | -0.518680 | 0.604066 | 0.787912 |
| MTOR | 0.046233 | -0.057040 | 0.479503 | 0.631657 | 0.789572 |
| MYC | 0.036033 | -0.053670 | 0.380451 | 0.703670 | 0.844404 |
| CASP3 | 0.027063 | -0.224020 | 0.299306 | 0.764752 | 0.850481 |
| TWIST1 | 0.030437 | -0.058700 | 0.298414 | 0.765433 | 0.850481 |
| MKI67 | -0.014180 | -0.011680 | -0.140220 | 0.888508 | 0.951973 |
| PTPRC | -0.00288 | -0.14908 | -0.03257 | 0.974025 | 0.987457 |
| ESR1 | 0.000747 | 0.502018 | 0.015724 | 0.987457 | 0.987457 |
